# WinnerTakeAlmostAll: A parsimony approach integrating multi-targets improves marker gene profiling

**DOI:** 10.1101/575100

**Authors:** Zewei Song, Xiaohuan Sun, Zhongkui Xia, Yangyang Jia, Zhu Jie, Chao Fang, Huanming Yang, Jian Wang

**Author notes:** Correspondence should be addressed to: Zewei Song.

## Abstract

Amplicon sequencing need to go from using single region to multiple region, one for higher resolution, one for technical replicates. WinnerTakeAlmostAll is an algorithm to calculate a minimum set of references based on the reference-query graph. We showed that this algorithm is superior in resolving species, as well as average out the run effect using technical replicates.

---

High throughput sequencing of the marker genes (i.e. amplicon sequencing) has become an effective tool for studying microbial communities. The essence of amplicon sequencing is to de-complicate the data, so that sequences with same origin were grouped together and counted for a “taxonomic unit” in the profile. These groups are named as “operational taxonomic units, OTUs” if using a similarity threshold in De novo or closed-ref methods^1^, or “sequence variants, SVs” if inferred from a self-derived error profile^2^. All current applications can only be applied to a single region (e.g. V4 of 16S for prokaryotes, ITS1 of ITS for fungi). Although researchers generated numerous data using a single variable region for profiling, it does not mean that the debate over the best region is over (). On the contrary, with the increasing discovery of new species, full-length ribosomal RNA needs to be weighed in for improved resolution^3^. Current long-read techniques can provide enough read length, but not enough throughput in the near future^4^. An alternative is to sequence multiple targets to represent the full-length gene using short read. But, we need a novel bioinformatic tools that can perform the “combine” function for these independent sequencing events. The ability to “average” technical replicates data is another promising application for the coming higher throughput. Increasing evidences has shown that run biases are prevalent in current sequencing platform^5,6^. We have showed that technical replicates were useful in detecting such biases. At the dawn of triplicating every sample, we need to prepare a new tool for such need.

We developed a new algorithm – winnerTakeAlmostAll (WTA) - that consider the alignment space (all alignment possibilities between query and reference) of multiple targets for a weighted profile. This method is inspired by the CAPITALIST mode in the unpublished aligning tool Burst^7^, as well as in earlier RNASeq analysis^8^, both of which were applied to a single sequencing event. The algorithm was embodied in our analysis package metaSeq wrapping using a snakemake pipeline^9^. All codes and scripts are open source and hosted at public repository^10^.

WTA approximates a minimum set of references by reducing the Query-Reference graph (QR graph). The QR graph is generated using the alignments considering multiple possibilities, either all hits above the threshold, all best hits, or any hit meet the users’ criteria. For experiment amplifies one variable region (target), this results in a graph with two types of nodes – queries and references – hereby referred as ref-node and query-node. Each query-node has one or more edges to ref-nodes. Each ref-node has one or more edges from query-nodes. There is no direct edge between query-nodes, or between ref-nodes. Instead, all query-nodes or ref-nodes can only reach another query-node or ref-node through another ref-node or query-node.

We then extended the QR graph to independent sequencing events (i.e. different region, same region but different runs). The resulting QR graph has n types of queries, in which n equals to the number of sequencing events (e.g. Figure 1a). A critical feature of QR graph is that it has a fix set of ref-nodes, and multiple types of query-nodes. This could be equal to the records in SILVA 16S database (as ref-nodes) and all sequenced variable regions (each as a type of query-nodes). Figure 1a is an example of the V1-V2 and V4 region of HMP 276D aligned to SILVA 16S database.

**Figure 1.**
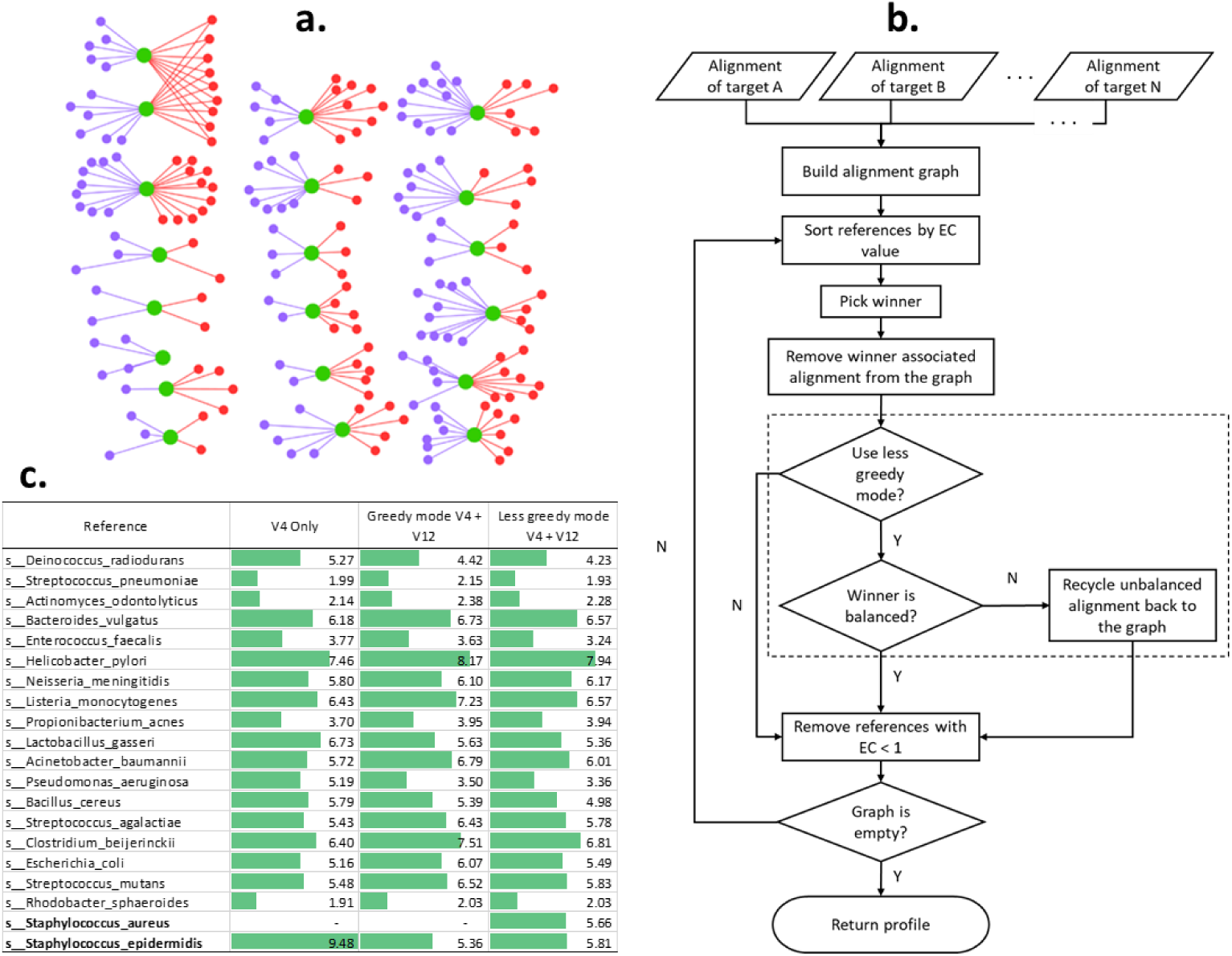
QR-graph and associated WTA flowchart and species abundance resolved using WTA compared to traditional alignment-based method. **a.** A QR graph using HMP-276D as example. Green nodes are the 20 bacteria species, aka ref-nodes. In the upper left corner are *Staphylococcus capitis* and *Staphylococcus epidermidis*. Purple nodes are queries sequenced using V1-V2 region at 2014, and red nodes are queries sequenced using V4 region at 2016, aka query-nodes. Edges represent eligible alignments. The two *Staphylococcus* species shared all V4 query-nodes, which will be resolved in the WTA algorithm. **b.** Flow chart of winnerTakeAlmostAll method. The recycle mechanism, aka less greedy mode is in the dashed box. **c.** Community profiles using V4 region only or combined with V1-V2 region using two modes of WTA.

After creating the QR-graph, WTA algorithm goes through an iteration until the QR-graph is empty. During each iterative step, a ref-node is picked as winner, and all queries connect to it are marked as discard (Figure 1b). For single target QR-graph, the ref-node with the most degree (i.e. the number of queries) is the winner. For multiple targets QR-graph, in which every ref-node has an array of abundance of connected queries, we proposed three methods:

1) Effective count (EC) = AVE – STDEV

2) Average

3) Medium

The effective count method takes into consider the balance among sequencing events. Unequal abundance among sequencing events get penalty against the average abundance. We assume that such imbalance originates from the redundant alignments in that targets (i.e. the query can alignment to more than one reference), partial of the alignment count should be assigned to another reference. In the case that we cannot resolve the overlapped ref-nodes, we give a penalty to the winner ref-node.

Similar to the metaphor in Burst (CAPITALIST mode)^7^, We can mimic our algorithm to a free market without regulation. Consider a market with multiple capitalists (references) who compete for profits (queries). In the first round of competition, the capitalist who owns the most profits will deprive them from the other capitalists, then become the winner. Some of the capitalists now owns zero profit, thus are kicked out. The competition continues until there is no profit and capitalist left in the market (i.e. the free market turns to a monopoly market). One useful addition to this mimic is that when there is more than one type of profit (i.e. multiple targets), the winner capitalist can maximize the profit by discard/recycle some of the profit back to the market. For example, a ref-node with abundance array [100, 50] has an expected EC of 39.6, but if it only keeps [50, 50], the expected profit increases to 50, which is a better solution for the winner. We thus introduce such recycle mechanism in our algorithm as the “less greedy” mode, comparing to the default “greedy” mode (Figure 2 in the dashed box). It has to be noticed that the less greedy mode is only compatible with the EC method, and is logically invalid for average and medium method.

**Figure 2.**
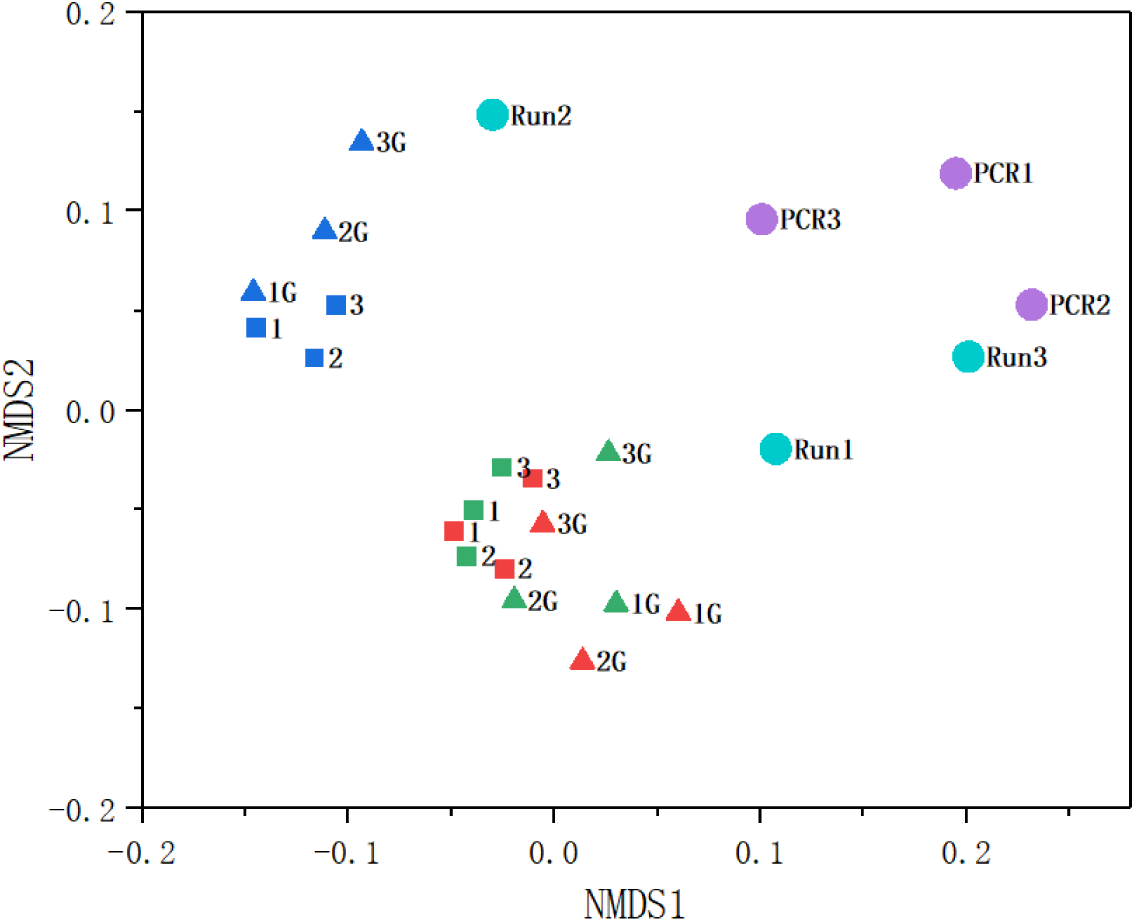
NMDS plots shows individual technical replicates versus “averaged” samples using WTA across either PCR (tan) or Run (purple). run1(green), run2(red), run3(blue), Combined three PCR (big cycle tan), combined three run (purple)

The entire algorithm is achieved under Python 3.6 using networkx package^11^ to achieve the graph operations. The aligner we recommend is Burst, but any aligner that is able to output multiple alignments can be used.

In order to bench marking our algorithms, we present two cases. In the first case, HMP 276D was sequenced on V1-V2 and V4 regions of bacterial 16S by two groups at 2014 and 2016, independently^12,13^. The two *Staphylococcus* species are identical at their V4 region but are not at V1-V2 region. The mock community was best resolved using the less greedy mode combining V1-V2 and V4 regions (Figure 1c). Aligning only V4 region to the references yielded similar result as the de novo clustering^13^. In the profile, one random *Staphylococcus* species got abundance around 10% (twice as expected). When considering both regions in the greedy mode, still only one species was resolved, but its abundance reduced to around 5% due to the penalty of imbalance. After switching to the less greedy mode, both species appeared in the profile with abundance around 5%. It is obvious that the recycle mechanism in our algorithm is critical for “combining” the multiple variable regions. Since the resolving power of the variable regions is taxonomic dependent, we argue that an alternative to choosing an “optimal” region is to sequence as many regions as possible. With our algorithms, the biases originate from one region can be compensate by “combing” multiple regions.

In the second case, we demonstrated how technical replicates can be “averaged” to reduce the run biases. We tested a dataset that was previously showed to have run biases^5^. In this dataset, three samples each had three PCR libraries, and each PCR library was sequenced in three runs. We saw a significant run effect, that data in run2 is significantly different than the other two runs, if de novo clustered the sequences. Here, we use the greedy mode to “average” out the three replicates from three runs or three PCR reactions. When average across runs, the resulting samples has the least variation in the NMDS plot (Figure 2). Here we choose the greedy mode since all potential OTUs were considered to be present in all samples, so if one references node is imbalanced, it is considered as a penalty to the run biases and can be compensated by “average” with other samples. We have seen that many OTUs previously identified at “variable” were have higher abundance in the new profile (data not shown).

Overall, WTA is the first parsimony algorithm that can utilize multiple targets in amplicon sequencing for a comprehensive profile. Our algorithm solves the dilemma of choosing the variable region in amplicon sequencing, also provide a way to “average” technical replicates, enabling the higher throughput processing of samples. WTA only need to process one sample at a time, so there is neglectable constrain on required memory (around three times of the raw sequence size). Computing time is linear with the sample size. Future implementation includes adding the weight of the similarities, reporting graph features, and use the QR-graph itself as a quantification measure.

## Acknowledgement

We would like to thank the helpful discussion with Dr. Al-Ghalith on the CAPITALIST idea.

## Online Methods

### Data prepare

For case I, we retrieved the data of Salipante et al. 2014 and Gohl et al. 2016 from SRA using the accession number SRP040453 and SRP069981. For the Salipante et al. 2014 dataset, we picked the one sequenced using MiSeq. For the Gohl et al. 2016 dataset, 276D were sequenced using multiple condition. We choose the first one out of the three technical replicates that was amplified by KAPA in the standard cycle time and DNA input.

For case II, we used the data in Song et al. 2018 archived in the Data Repository for the University of Minnesota (DRUM, https://conservancy.umn.edu/). Accession number for run1 is 190418, and for run2 and run3 is 190417. In total we used 27 sequence samples (3 samples X 3 PCR libraries X 3 runs). All samples were sequenced at the University of Minnesota Genomic Center following the protocol of Gohl et al. 2016.

All sequence files were passed through quality following our standard pipeline (Song et al. 2018, see also https://github.com/ZeweiSong/FAST). In brief, corresponding primers were trimmed, paired sequenced were merged, then sequences with maximum expected error > 1 were discarded.

### Profile interpretation

Processed sequences were aligned to reference database, SILVA (version, bacteria 16S) for case I and UNITE (version, fungal ITS) for case II. Sequences were aligned to reference using Burst. For case I, ALLPATHS mode was used to find all best hits. In case II, FORAGE mode was used to find all hits above the 0.97 identity.

### WTA profile

For case I, we generated profiles using three sets of parameters:

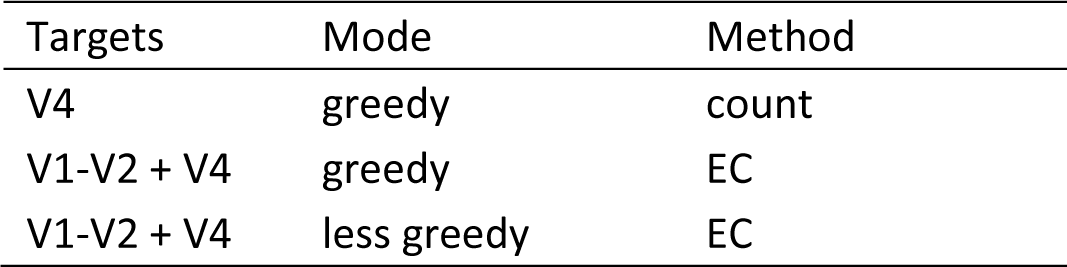

Since some species has multiple records, we collapse those after the alignment by changing the replacing the target name with common names.

For case II, we generated WTA profile using greedy mode and average method. We first generated the 27 profile using single sequence. Then PCR replicates and run replicates profile were generated. NMDS plot was calculated using the vegan package in R.

